# Genome-based comparison between the recombinant SARS-CoV-2 XBB and its parental lineages

**DOI:** 10.1101/2022.12.20.521197

**Authors:** Fabio Scarpa, Daria Sanna, Ilenia Azzena, Marco Casu, Piero Cossu, Pier Luigi Fiori, Domenico Benvenuto, Elena Imperia, Marta Giovanetti, Giancarlo Ceccarelli, Roberto Cauda, Antonio Cassone, Stefano Pascarella, Massimo Ciccozzi

## Abstract

Recombination is the main contributor to RNA virus evolution, and SARS-CoV-2 during the pandemic produced several recombinants. The most recent SARS-CoV-2 recombinant is the lineage labeled XBB, also known as Gryphon, which arose from BJ.1 and BM. 1.1.1. Here we performed a genome-based survey aimed to compare the new recombinant with its parental lineages that never became dominant. Genetic analyses indicated that the recombinant XBB and its first descendant XBB.1 show an evolutionary condition typical of an evolutionary blind background with no further epidemiologically relevant descendant. Genetic variability and expansion capabilities are slightly higher than parental lineages. Bayesian Skyline Plot indicates that XBB reached its plateau around October 6, 2022 and after an initial rapid growth the viral population size did not further expand, and around November 10, 2022 its levels of genetic variability decreased. Simultaneously with the reduction of the XBB population size, an increase of the genetic variability of its first sub-lineage XBB.1 occurred, that in turn reached the plateau around November 9, 2022 showing a kind of vicariance with its direct progenitors. Structure analysis indicates that the affinity for ACE2 surface in XBB/XBB.1 RBDs is weaker than for BA.2 RBD. In conclusion, nowadays XBB and XBB.1 do not show evidence about a particular danger or high expansion capability. Genome-based monitoring must continue uninterrupted in order to individuate if further mutations can make XBB more dangerous or generate new subvariants with different expansion capability.

## 1. Introduction

Nowadays, the world is being confronted with the ongoing current pandemic of COVID-19 caused by the SARS-CoV2 identified for the first time in December 2019 during a pneumonia outbreak in Wuhan (China) [1]. After the first cases, it rapidly emerged as a worldwide concern [2]. On 11 March 2020, with a total of 149,295 confirmed cases, the World Health Organization (WHO) declared COVID-19 a pandemic (https://www.who.int/director-general/speeches/detail/who-director-general-s-opening-remarks-at-the-media-briefing-on-covid-19---11-march-2020). In December 2022, WHO declared that, over 645 million confirmed cases and over 6.6 million deaths have been reported globally (https://www.who.int/publications/m/item/weekly-epidemiological-update-on-covid-19---14-december-2022).

Due to the continuing evolution of SARS-CoV-2 and the expected generation of new variants, SARS-CoV-2 infections are likely to remain a problem for the time being in most countries [3]. Indeed, SARS-CoV-2 is a positive-sense single-stranded RNA virus, with a high error rate in RNA replication thus mutation and evolutionary fitness mostly affecting its transmissibility [4].

Accordingly, during the pandemic SARS-CoV-2 significantly mutated over the time producing many lineages and sub-lineages with different expansion capabilities [5]. For the generation of new viral variants, recombination events may play a non-negligible role. Indeed, recombination is generally the main contributor to RNA virus evolution [6], as well as the re-assortment which, however, is present only in RNA viruses with segmented genomes such as the influenza virus (see e.g. Mugosa et al [7]). Of course, recombination within different lineages requires the co-circulation and the co-infection of the viruses in the same host [8]. As for the newly discovered variants, the occurrence recombinants must be monitored and requires a constant surveillance. The most recent SARS-CoV-2 recombinant is the lineage labeled as XBB, which has been also nicknamed Gryphon [9]. XBB lineage is a recombinant of BJ.1 (also known as *Argus*) and BM.1.1.1 (also known as *Mimas*), both belonging to the BA.2 lineage (https://www.who.int/activities/tracking-SARS-CoV-2-variants), with the following Spike mutations in addition to those typical of its progenitor lineage: V83A, Y144-, H146Q, Q183E, V213E, G252V, G339H, R346T, L368I, V445P, G446S, N460K, F486S, F490S, and T11A (E), K47R (ORF1a), G662S (ORF1b), S959P (ORF1b) and G8 (ORF8) as additional ones outside of the spike protein (https://outbreak.info/compare-lineages) [10].

As of December 2, 2022, from the sequences submitted to GSAID, XBB and its descendant showed a global sequence prevalence of around 7% (https://gisaid.org/phylodynamics/global/nextstrain/). The Technical Advisory Group on SARS-CoV-2 Virus Evolution (TAG-VE) discussed on the growth advantage of this sublineage and some early evidence on clinical severity and reinfection risk in several countries (i.a. Singapore and India) (https://www.who.int/news/item/27-10-2022-tag-ve-statement-on-omicron-sublineages-bq.1-and-xbb). Recently, an alarming evidence has been provided on the capacity by XBB and XBB.1 of evading antibody-mediated immunity conferred by vaccination or previous infection although fully vaccinated subjects remain still protected from hospitalization and death [11]. For these strongly immunoevasive properties, XBB require a constant genome-based epidemiological surveillance and clinical monitoring of disease characteristics.

In such a context, here we performed a genome-based survey focused on genetic variability/phylodynamics and structural analyses in order to obtain an as complete as possible assessment of the evolutionary potential for epidemiological trajectory and dangers that XBB and its descendant XBB.1 could inflict to the population. In particular, the recombinant XBB and its first descendant XBB.1 have been compared with its parental lineages, in order to understand if these two lineages present new biological features that may candidate them to spread quickly and possibly outcompete the parental lineages.

## 2. Materials and Methods

In order to locate XXB and XXB.1 into an evolutionary perspective, genomic epidemiology of SARS-CoV-2 Omicron variants has been reconstructed by using a subsampling focused globally over the past 6 months, built with nextstrain/ncov (https://github.com/nextstrain/ncov) (available at https://gisaid.org/phylodynamics/global/nextstrain/), including all genomes belonging to the GSAID Clade 21 L (Omicron) (2371 of 2946 genomes sampled between January 2022 and December 2022).

After the first genomic assessment, in order to perform a genetic comparison between XXB and XBB.1 and their parental lineages, four subsets were build: BJ.1 (n= 134), BM.1.1.1 (n= 70) XBB (n= 1,602) and (n= 1,841) and for each dataset independently the above described genetic analyses were performed.

Genomes were aligned by using the algorithm L-INS-I implemented in Mafft 7.471 [12], producing datasets of 29,729 (BJ.1), 29,719 (BM.1.1.1), 29, 717 (XBB) and 29,725 (XBB.1) bp long. Manual editing was performed by using the software Unipro UGENE v.35 [13]. The software jModeltest 2.1.1 [14] was used to find the best probabilistic model of genome evolution with a maximum likelihood optimized search. Times of the most recent common ancestor and evolutionary rate were estimated by using Bayesian Inference (BI), which was carried out using the software BEAST 1.10.4 [15] with runs of 200 million generations under several demographic and clock model. In order to inference on the best representative output the selection of the better model was performed by the test of Bayes Factor [16] by comparing the 2lnBF of the marginal likelihoods values following Mugosa et al. [7]. The software Beast was also used to draw the Bayesian Skyline Plot (BSP) and lineages thorough times for XBB, XBB.1 BJ.1 and BM. 1.1.1 with runs of 200 million generations under the Bayesian Skyline Model with the uncorrelated log-normal relaxed clock model.

All datasets were built by downloading genomes form GSAID portal (https://gisaid.org/) available at November 16, 2022.

Break-point of the recombination event was individuated on a dataset sequence composed by all investigated genomes belong to the two parental lineages and the recombinant linage (BJ.1+BM.1.1.1+XBB).

The mutations characterizing the XBB and XBB.1 SARS-CoV-2 Spike lineages were individuated by using consensus sequences obtained applying a cutoff of 75% sequence prevalence on all available sequences. After individuation, mutations were verified by comparing results of the "Lineage Comparison" web page of GISAID. Homology models of the mutant Spikes were created by means of the software Modeller 10.3 [17]. Spike structures were displayed and analyzed with the graphic program PyMOL [18]. Surface electrostatic potential has been calculated with the program APBS [19] and displayed as a two-dimensional projection with the SURFMAP software [20]. SURFMAP implements a method of "molecular cartography" by means of which a protein three-dimensional surface can be projected onto a two-dimensional plane. Distribution of physico-chemical features over the protein surface can be analyzed and compared using the two-dimensional map. Net charge were calculated by the software PROPKA3 [21]. Foldx5 [22] was applied to optimize the side chain conformation of the models built by Modeller using the Foldx5 function "RepairPDB". Interaction energy between the Spike RBD and ACE2 were predicted with the Foldx5 function "AnalyseComplex". Interface residue-residue interactions were assessed with the Foldx5 "PrintNetwork" function. In silico mutagenesis was obtained with the built-in functions available within PyMOL. In silico alanine scanning of the residues at the interface between RBD and ACE2 was carried out using the method available via the web server DrugScore(PPI) [23]. The method is a fast and accurate computational approach to predict changes in the binding free energy when each residue at the subunit interface is mutated into alanine. The predicted net charge of the domains at pH=7.0 (set as a reference pH, though not necessarily reflecting the physiological environment) was calculated by means of the program PROPKA3. The predicted overall net charge is dependent on the combination of the sidechain charges. The net charge is influenced by the interactions of the side chains with the surrounding atoms. To sample the variability of the interactions, 100 homology models of each RBD have been calculated with Modeler. Indeed, the Modeler refinement stage of the homology modelling can produce models differing for the conformation of side chains. Each model has been optimized by the Foldx5 "RepairPDB" procedure and the net charge has been calculated by PROPKA3. The final charge was the average of the 100 charges and the variability estimated by the standard error. To sample the fluctuations of the side chain conformations, the same procedure was applied to estimate the interaction energy of the complex ACE2-RBD.

## 3. Results

Phylogenomic reconstruction (Fig. 1) indicates that XBB and XBB.1 (GSAID Clade 22F) genomes clustered within the not-monophyletic GSAID Clade 21L. More specifically, and as expected, they are evolutionary close to genomes of BA.2 which represent their ancient progenitor. Results of the Bayes Factor on the four datasets revealed that the Bayesian Skyline Model under the lognormal uncorrelated relaxed clock model fitted data significantly better than other tested demographic and clock models for all analyses datasets.

**Figure 1.**
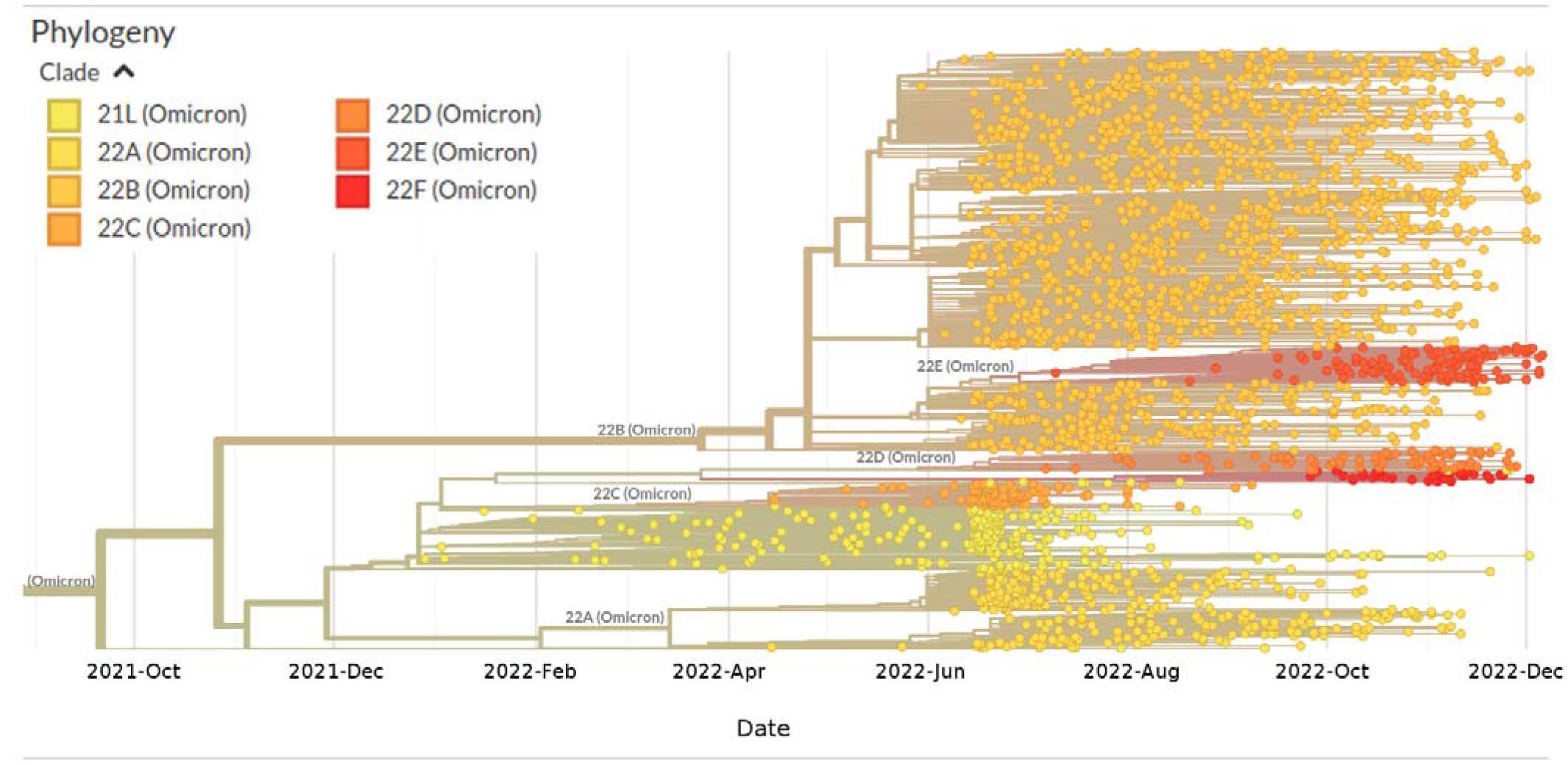
Highlight of the Omicron Clade (GSAID Clade 21L) in the time-scaled phylogenetic tree of a representative global subsample of 2371 SARS-CoV-2 genomes sampled between January 2022 and December 2022. Figures has been edited by using the software GIMP 2.8 (available at https://www.gimp.org/downloads/oldstable/).

The Time of the Most Recent Common Ancestor (TMRCA) of the clade composed by XBB+XBB.1 is placed 115 days before November 12, 2022, i.e. July 20, 2022, with a date interval confidence of 176 - 68 days (i.e. May 20 - September 5).

Bayesian Skyline Plot (BSP) of the parental lineage BJ.1 (Fig. 2a) showed that viral population has undergone an increase in size starting from about 110 days before October 28, 2022 (i.e. July 10, 2022), reaching the peak about 73 days before October 28, 2022 (i.e. August 16, 2022). Lineages through times plot (Fig. 2b) indicates that the maximum number of lineages has been achieved around 73 days before October 28, 2022 (i.e. August 16, 2022).

**Figure 2.**
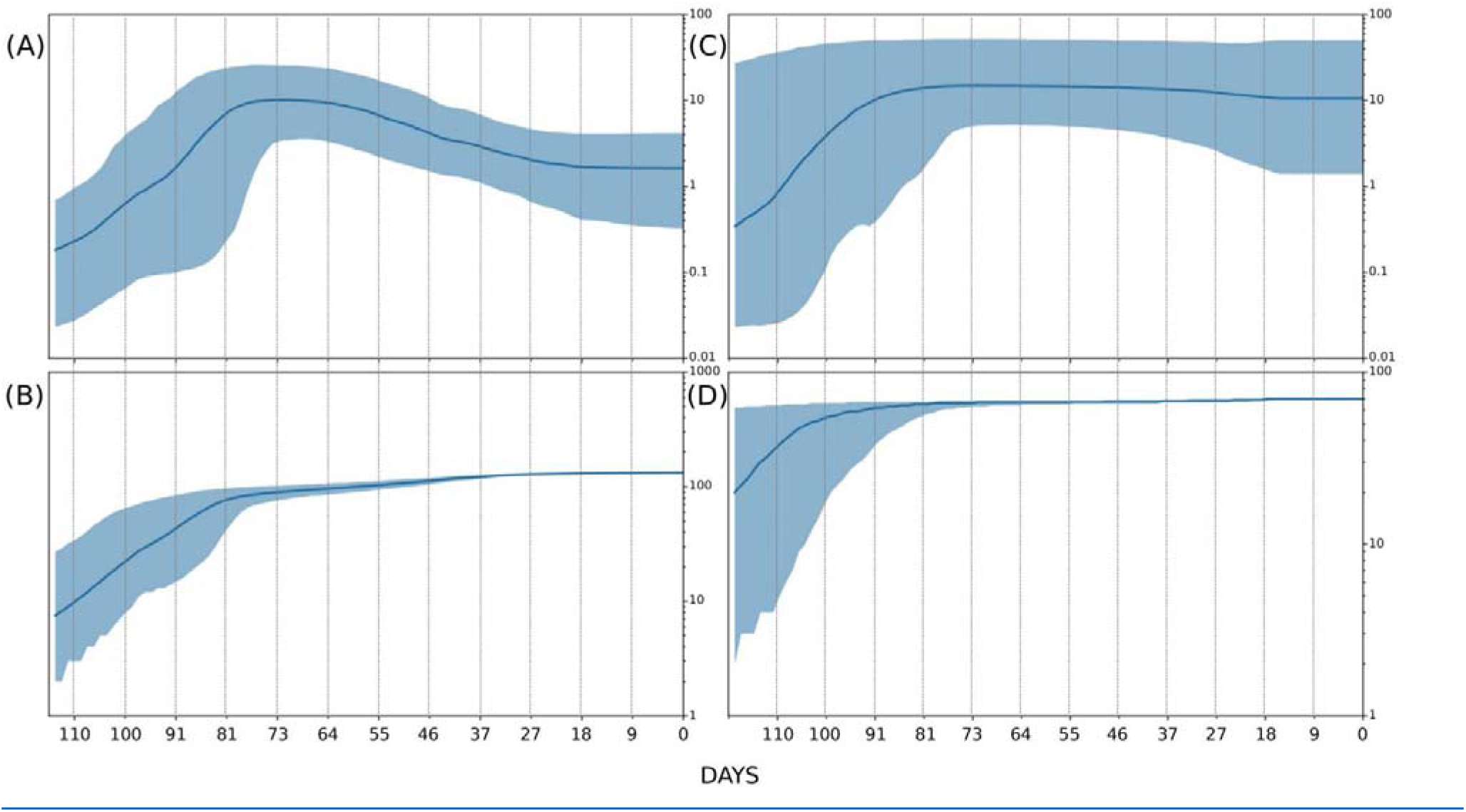
Bayesian skyline plot and lineages through time of SARS-CoV-2 BJ.1 (A, B) and BM.1.1.1 (C, D) variants. The viral effective population size (A, C) and the number of lineages (B, D) in the y-axis are shown as a function of days (x-axis).

BSP of the parental lineage BM.1.1.1 (Fig. 2c) indicated that after the initial expansion in population size, the peak was reached around 91 days before November 3, 2022 (i.e. August 4, 2022) when started the plateau. Lineages through times plot (Fig. 2d) indicates that the maximum number of lineages has been achieved around 100 days before November 3, 2022 (i.e. July 25, 2022).

BSP of the recombinant XBB (Fig. 3a) showed that after a first period of flattened genetic variability, the viral population has undergone an increase in size starting from 46 days before November 12, 2022 (i.e. September 27, 2022), reaching the peak around 10 days after (i.e. October 6, 2022) and followed by a plateau phase that lasted until few days before November 12, 2022 when a reduction in viral population size occurred. Lineages through times plot (Fig. 3b) indicates a moderately slow increase of the number of lineages until around 46 days before November 12, 2022 (i.e. September 27, 2022), than the number of lineages stopped its increasing.

**Figure 3.**
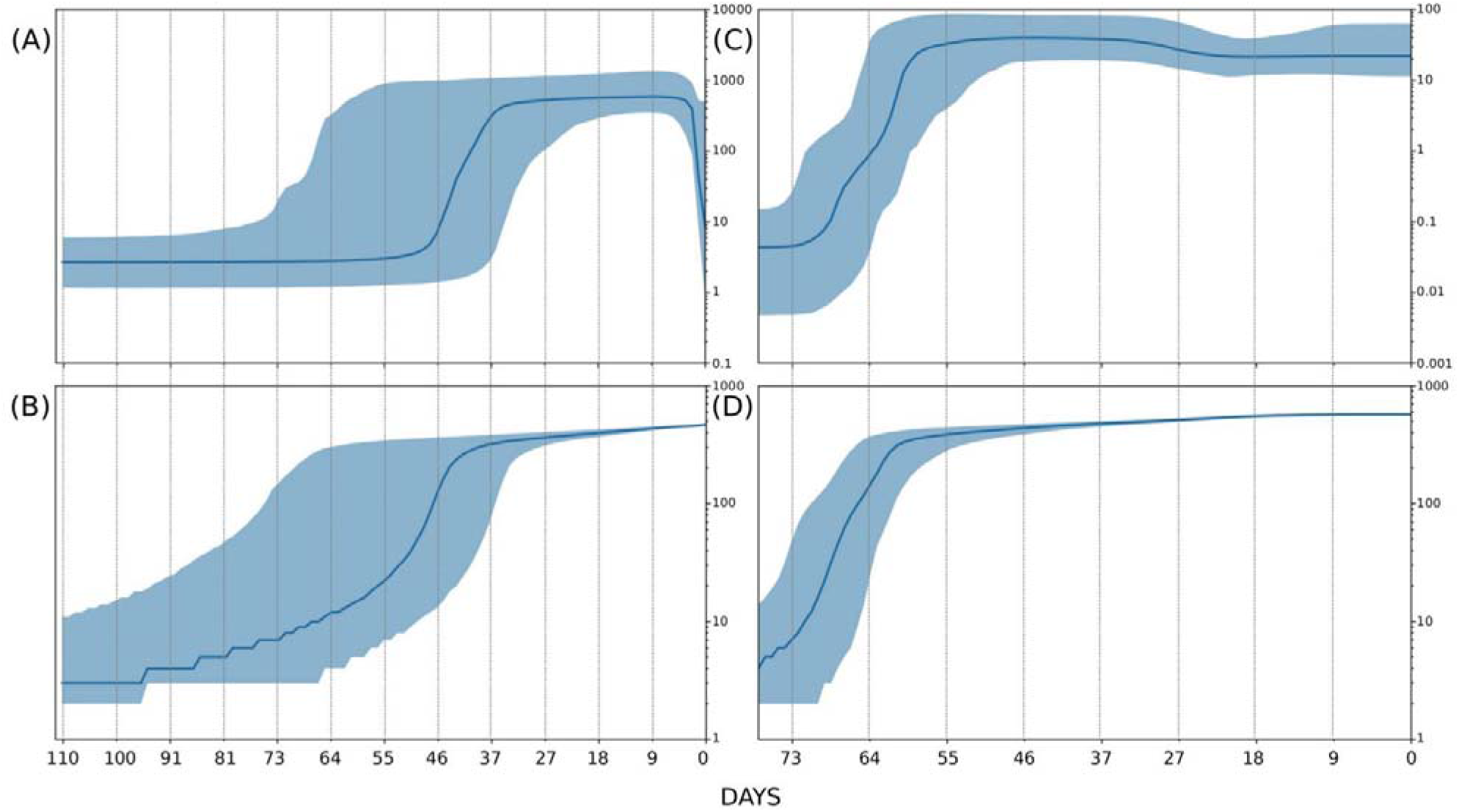
Bayesian skyline plot and lineages through time of SARS-CoV-2 XBB (A, B) and XBB.1 (C, D) recombinant. The viral effective population size (A, C) and the number of lineages (B, D) in the y-axis are shown as a function of days (x-axis).

BSP of the sublineage XBB.1 (Fig. 3c) showed a brief period of flattened genetic variability after that an increase of the viral population size started around 70 days before November 9, 2022 (i.e. August 31, 2022) reaching its peak around 60 days before November 9, 2022 (i.e. September 10, 2022), then the plateau which is still ongoing without variation in genetic variability and viral population size. Lineages through times plot (Fig. 3d) indicates a moderately slow increase of the number of lineages until around 64 days before November 9, 2022 (i.e. September 6, 2022), than the number of lineages stopped its increasing.

Evolutionary rate of the four tested lineages amount to 1.4×10^-3^ [95% HPD 6.4×10^-4^ - 2.4×10^-3^], 1.3×10^-3^ [95% HPD 7.4×10^-4^ - 1.9×10^-3^], 7.6×10^-5^ [95% HPD 4.57×10^-5^ - 1.0×10^-4^] and 6.3×10^-4^ [95% HPD 5.2×10^-4^ - 7.3×10^-4^] subs/sites/years for BJ.1, BM.1.1.1, XBB and XBB.1, respectively.

The structural analysis has been restricted to the most widespread lineages XBB and XBB.1 for which a wealth of sequential information is available in the databanks that allows a more precise definition of mutations prevalence. The N-terminal domains (NTD) of the two variant Spikes differ for the mutation G252V that is present in XBB.1 and is missing in XBB. On the contrary, the two RDB domains share an identical mutational profile. The structural properties of the two variant Spikes have been compared to those of the parent BA.2 Spike.

The mutation V83A occurs in a site within an exposed loop (Fig. 4). The deletion of Y144 and the substitution H146Q are in the β-strand encompassed by the sequence positions 140-146. The two mutation positions are near the putative binding site of the AXL receptor shown in Fig. 4 [24,25]. The mutation G152V is specific of XBB.1 and it occurs within a loop also putatively involved in the interaction with the AXL receptor. The mutation Q183E occurs in an exposed loop and introduces a negatively charged residue that decreases the overall net charge and alters the distribution of the surface electrostatic potential (Table 1) with respect to the parental BA.2.

**Figure 4.**
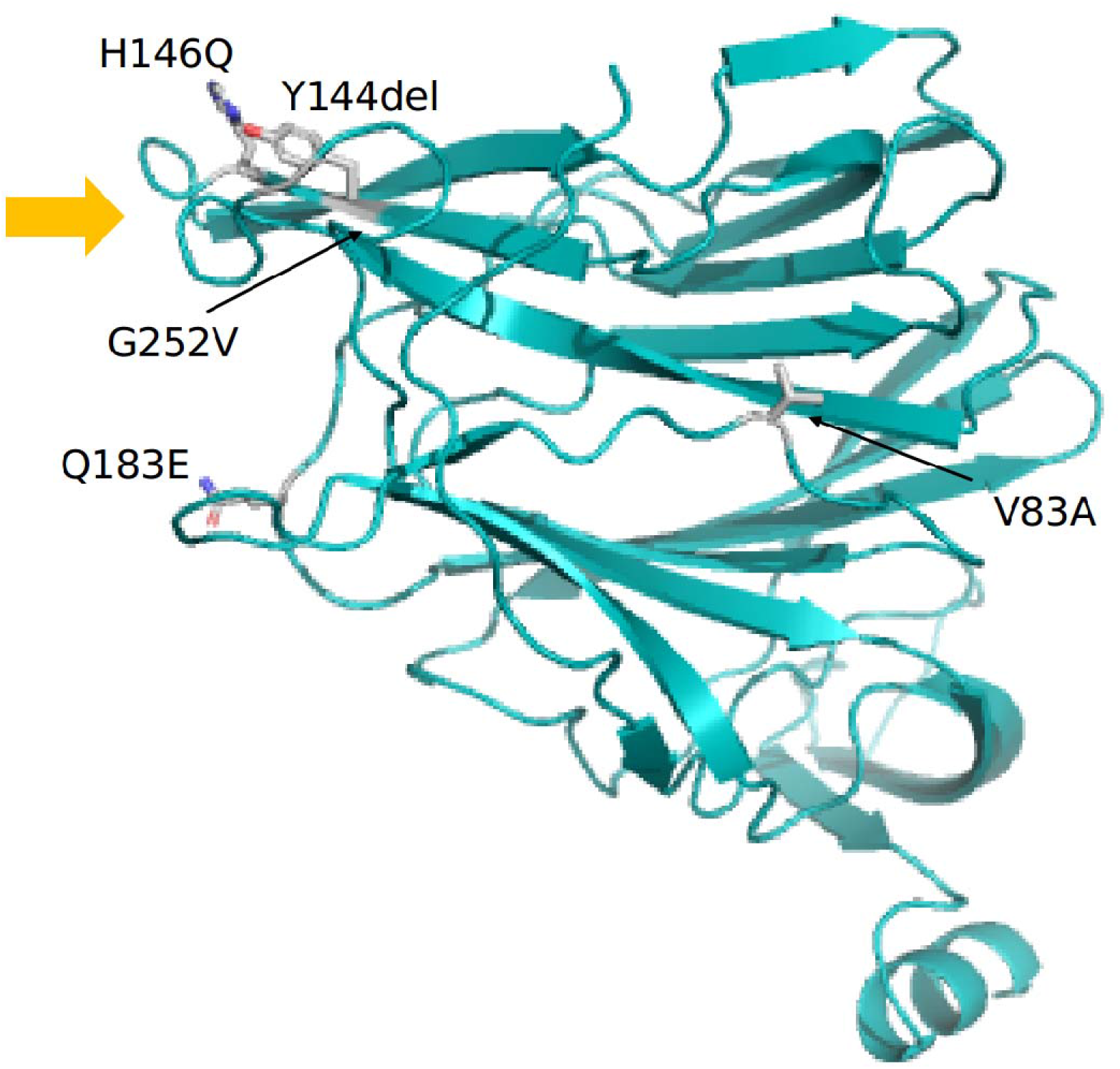
The N-terminal domain represented as a cartoon model. The mutated sites specific of XBB or XBB.1 are indicated by gray sidechains reported as labeled stick models. Orange arrow marks the domain portion putatively involved in interaction with AXL.

**Table 1.** Nextstrain clade, Pango lineage and WHO labels of the investigate lineages showed in Figure 1.

| Nextstrain Clade | Pango Lineage | WHO label |
| --- | --- | --- |
| 21L (Omicron) | BA.2 | o (Omicron) |
| 22A (Omicron) | BA.4 | o (Omicron) |
| 22B (Omicron) | BA.5 | o (Omicron) |
| 22C (Omicron) | BA.2.12.1 | o (Omicron) |
| 22D (Omicron) | BA.2.75 | o (Omicron) |
| 22E (Omicron) | BQ.1 | o (Omicron) |
| 22F (Omicron) | XBB | o (Omicron) |

**Table 2.** Net charge of RBD and NTD domains and interactions energy in the complex ACE2-RBD.

| Variants | RBD<sup>a</sup><br>net charge | NTD<br>net charge | RBD-ACE2<sup>a</sup><br>(Kcal/mol) |
| --- | --- | --- | --- |
| BA.2 | 5.19 ± 0.01 | 0.95 ± 0.04 | -6.19 ± 0.32 |
| XBB | 5.45 ± 0.02 | -1.24 ± 0.04 | -3.54 ± 0.30 |
| XBB.1 | = | -1.18 ± 0.03 | = |

<sup>a</sup> “=” means identical value

The XBB and XBB.1 RBD domains share the same mutation pattern. The mutation R346T occurs in a solvent exposed loop while L368I is within a short helix between positions 365-371 (Fig. 5). The mutation V445P is in a loop near the interface to ACE2 although the side chain is not in direct contact with any ACE2 residue. On the contrary, G446S and F486S are in the ACE2 interface. The mutations N460K and F490S are also not directly involved in the ACE2 interface. Of note, BA.2 shows the mutation Q493S that is missing in XBB and XBB.1, in a site that is within the ACE2 interface.

**Figure 5.**
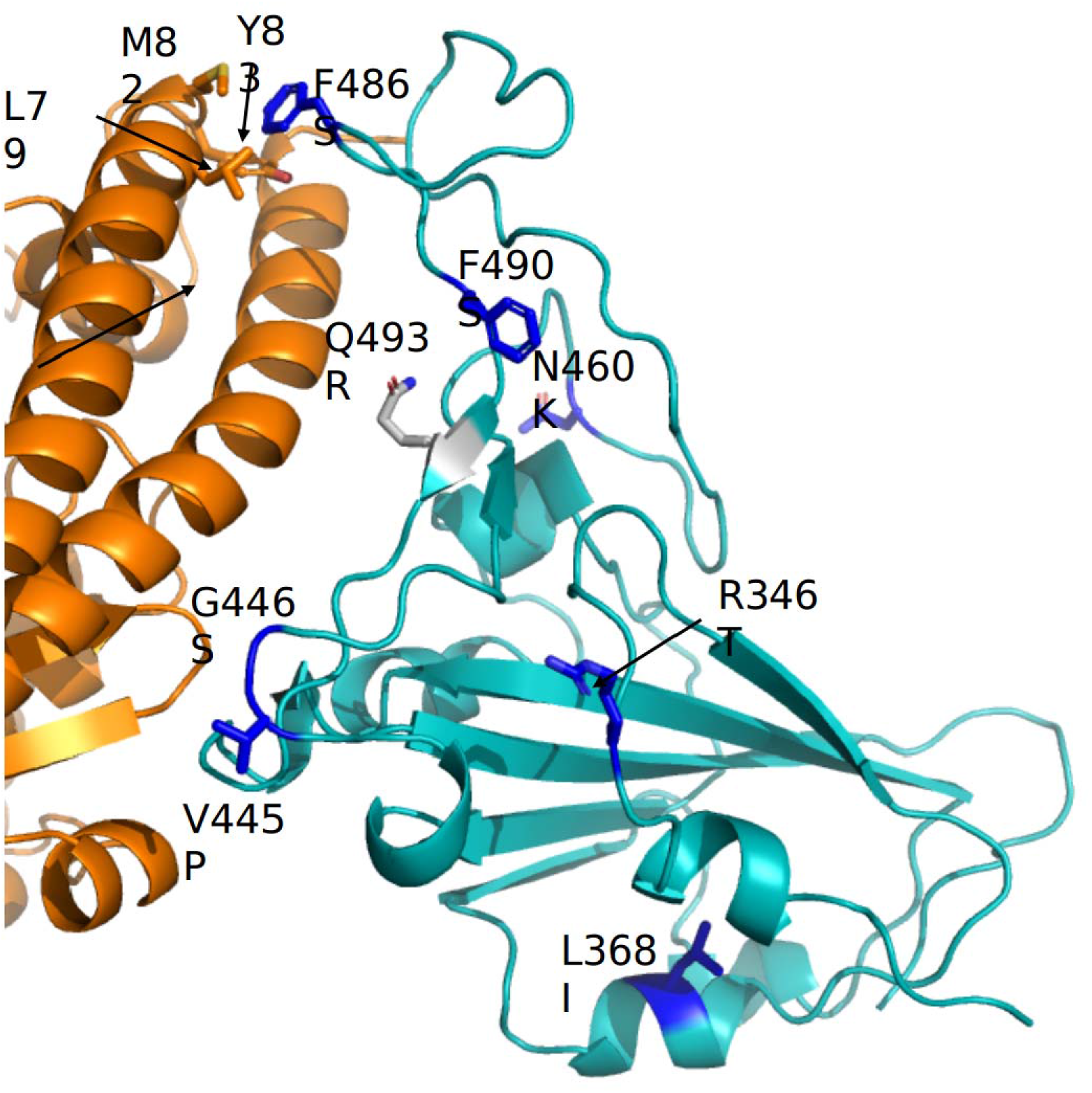
Portion of the complex between ACE2 (orange cartoon) and Spike RBD (teal cartoon). The XBB/XBB.1 specific mutated sites are highlighted with blue stick side chains labeled according to the corresponding mutation. The gray side chain marks the mutation specific to BA.2. Ace2 residues interacting with F486 are displayed as orange sticks.

The net charges of the BA.2 and XBB/XBB.1 are comparable although the positive surface potential distribution differs in the two cases. In fact, in the BA.2 RBD the positive potential seems to be more intense at the ACE2 interface area than in XBB/XBB.1. The prediction of the interaction energy between RBD and ACE2 suggests that BA2 RBD has a more stable interaction with this receptor than XBB/XBB.1 RBDs, within the accuracy limits of the predictive model. The putative weakening of the interactions may be in part explained by the XBB/XBB.1 mutations F498S. Indeed, alanine scanning carried out with the server DrugScore (PPI) predicts that F498 is an interface hot spot as its replacement with an Ala residue induces a loss of interaction energy of about 3.0 Kcal/mol. In the Wuhan RBD, F498 establishes hydrophobic interactions with the ACE2 residue L79, M82 and Y83.

## Discussion

The SARS-CoV-2 XBB recombinant is the most recent product of a recombination event occurred during the current COVID-19 pandemic. As all newly discovered variants, XBB requires a stringest evaluation of its genomic differences from its parental lineages in order to estimate its capabilities for expansion and contagiousness as well as its pathogenicity features, including immunoevasion. Here we sought for a deep insight into the evolutionary and structural patterns of the SARS-CoV-2 recombinat XBB and its first descendant XBB.1 by means a genome-based approach using of all genomes available in GSAID at November 16, 2022.

Phylogenomic reconstruction (Fig. 1) indicates that genomes of XBB and XBB.1 (GSAID Clade 22F) belong to a monophyletic group, which in turn fall within the wide and heterogeneous GSAID Clade 2 as their parental lineages BM. 1.1.1 and BJ.1. This is not surprising considering that they belong to the same sublineage (BA.2) and share a (not direct) common ancestor.

Similarly to what was recently observed for the variants BA.2.75 (nicknamed *Centaurus*) [26] and BQ.1 (nicknamed *Cerberus*) [27], the recombinant XBB and its first descendant XBB.1 show an evolutionary condition typical of an evolutionary blind background with no further epidemiologically relevant descendant that present features of concern.

In the near past the same condition was also observed in the BA.2.12.1 variant which indeed has not produced further new sub-lineages and whose genomic global sequence prevalence has been declining during time until its almost complete disappearance. Indeed, in the phylogenomic reconstruction, BA.2.12.1 variant (GSAID Clade 22C), BA.2.75 (GSAID Clade 22D), BQ.1 (GSAID Clade 22E) presented branches' length that suggested the lack of a rapid diversification [26, 27], similarly to the clade composed by XBB + XBB.1 in the current reconstruction, which does not highlight any features typical of an epidemiologically dangerous lineage at the beginning of its evolutionary path.

The common ancestor to all analyzed genomes of XBB is temporally placed 115 days before November 12, 2022 (which is most recent collection date), i.e. July 20, 2022. This molecular dating calibration predates of about one month (23 days) the first detected genome of XBB (for which a complete sampling date is available), which was isolated in India on August 13, 2022. Phylogenomic reconstruction set a genome of BJ.1 from Bangladesh (EPI_ISL_14970755) external to the clade of XBB and XBB.1 in a basal position, while other member of BJ.1 are placed within another sub-cluster. The position in the phylogenomic tree of the genome of BJ.1 from Bangladesh, together with the high genome prevalence in Eastern India and Bangladesh of both XBB and its parental lineages, suggest the lack of multiple introductions and the occurrence of one recombination event (probably occurred in a human) during the long branch phase, before the parental lineages were in broad circulation. During the recombination event the parental BJ.1 played the role of *Donor* while BM.1.1.1 is the *Acceptor* (*sensu* Focosi & Maggi [8]). The role of the parental lineages in the recombination event is further confirmed by the position of the breakpoint which is located in the genome between the nucleotidic position 22,901 and 22,939 in the SARS-CoV-2 reference genome (NC_045512.2), which correspond to the position 1,339 - 1,377 in the Spike protein gene sequence NC_045512.2 (21563..25384). More specifically the breakpoint occurred in the middle of the Receptor Binding Domain (around S:450-460), region for which the recombinant XBB carries the same mutation of BJ.1 until to G446S and from N640K carries the same mutations of BM. 1.1.1.

As of today the recombinant lineage XBB appear to be common in Asia, but according to dating results here proposed, XBB circulated undisturbed for about a month before being detected and for several months before causing concern. As recently pointed out for several variants newly generated (i.a. BA.2.75 and BQ.1) this is not the feature of a highly expansive variant [26,27], which typically explodes much faster in terms of numbers of infections and population size, as it occurred for the Omicron variant (B.1.1.529) for instance, which became predominant in a few time [28]. Indeed, BSP indicates an initial period of about 60 days with a flattened genetic variability and a very small viral population size. In accordance with the increase of number of lineages depicted by the lineages through times plot, XBB showed an increase of its genetic variability and a consequent increase in population size started around September 27, 2022. After a rapid growing of the viral population size, the peak was reached in about 10 days and the plateau phase has begun around October 6, 2022. The period of quick growth has been an evolutionary advantage for XBB, which has been able to replace its parental lineages BJ.1 and BM. 1.1.1 in Asia. However it should be pointed out that as of today, worldwide, it has been never expanded and around November 10, 2022 its population size is reduced as a consequence of a lower level of genetic variability. In that period it happened a kind of vicariance with its first sub-lineage XBB.1, which indeed showed an increase of the number of lineage and of the viral population size shifted forward a few days in time. Indeed, XBB.1 reached its plateau around November 9, 2022 after a moderate increase of the number of lineage and genetic variability. These replacements, both that of XBB vs its parents and that of XBB.1 vs XBB, have been possible just because of the lack of a sufficient strength of the variant allowing it to prevail over all others. BJ.1 and BM. 1.1.1 have never become dominant and their genome sequence prevalence was very low even before XBB arose. Notably, their evolutionary rate of 1.4×10^-3^ subs/sites/years for BJ.1 and 1.3×10^-3^ subs/sites/years for BM.1.1.1 are not so fast, as depicted by the BSP plots whose indicate that the peak phase has been reached around August 16, 2022 and July 25, 2022, respectively, with low level of increasing of lineages in both cases. This growth arrest of their population size allowed the increasing in genomic sequence prevalence of XBB although it has a lower evolutionary rate (7.6×10^-5^ subs/sites/years). On the contrary XBB.1 easily replaced its direct progenitor thanks to a faster evolutionary rate of 6.3 x 10^-4^ subs/sites/years which present a difference equal to a factor of 10^-1^.

The structural comparison between the three variants BA.2, XBB and XBB.1 suggests interesting differences. The NTDs of XBB and XBB.1 possess a more negative charge with respect to BA.2 that spreads over a large part of the molecular surface. Moreover, the characterizing mutations Y144del and H146Q occur near the putative site of interaction with the AXL receptor, that possesses an overall negative charge. In theory, these observations allow for the speculation that XBB and XBB.1 NTDs has a weaker propensity to interact with the AXL receptor and with the negatively charged sialosides displayed on the cell surface. Very likely, these changes also affect the interaction with the host immune system. Likewise, the mutations on XBB (identical in XBB.1) RBD do not significantly change the net charge that remains as much positive as in BA.2. However, the mutations alter the distribution of the positive electrostatic potential on the domain surface. In XBB/XBB.1 the positive potential becomes less localized at the ACE2 interface with respect to the BA.2 RBD. Moreover, the mutation F486S apparently destabilizes the complex ACE2-RBD. Once more, these observations suggest that XBB/XBB.1 RBDs may have a weaker affinity for ACE2 surface. Moreover, the alterations of the physico-chemical properties of Spike induced by mutations may potentially modify the interaction with the host immune system components.

Most of the mutations of the SARS-CoV-2 variants analyzed in this article fall in the NTD, RBD and RBM domains while most of the S2 region is highly conserved. This could probably enforce the hypothesis that NTD, RBD and RBM regions of the spike protein contain numerous strong B cell epitopes with the ability to readily elicit a strong neutralizing antibody response. Accordingly, their numerous mutations are considered to be largely responsible of the capacity of XBB and XBB,1 variants of massively evading neutralization by antibodies in both the vaccinated and the infected subjects as well [11]. Moreover, it has been suggested that Spike glycoprotein could conceal each of its immunodominant domains by adopting the closed conformation [29].

## Conclusions

In conclusion, the genome-based survey the SARS-CoV-2 recombinat XXB and on its first sublineage XBB.1 suggest that although this new variant presents several spike mutations of interest (and overall highly immune-evasive capacity to escape from neutralizing antibodies [30]), it currently does not show evidence about its particular dangerous or high expansion capability, although they are strongly immunoevasive. Recombination is a common evolutionary tool in the family Coronaviridae [31], as well as for RNA viruses in general [8]. As of now, XBB represents the first recombinant with showed a growth advantage that deserved to be investigated. Indeed, by becoming regionally dominant, it has caused concern,suggesting the need for an in-depth survey. Nonetheless, its expansion capability appears to be very limited with the peak reached in October 6 for XBB and November 9 for XBB.1. Data here reported indicates an initial quick growth followed by a long period of flattened genetic variability, very distant, in term of expansion capabilities, from an epidemiologically dangerous lineage as shown at the beginning of the pandemic where population size presented an extremely vertical curve (see, i.a., Lai et al. [32]).

The genome-based surveillance must continue for all SARS-CoV-2 lineages and variants in order to detect any new possible expansion. Structural interpretation of mutations provides a mean to formulate hypothesis for explaining and predicting the epidemiological behaviour of the variants. New further mutations can make XBB more dangerous or generate new subvariants and accordingly, in the near future the attention must be focused on its descendant, in order to verify their expansion capability and biological features for a better understanding of the pandemic.

## Acknowledgments

This research was funded from by FONDAZIONE DI SARDEGNA <u>bando 2022-2023</u> for the Dipartimento di Scienze Biomediche - UNISS (to Daria Sanna and Fabio Scarpa). Marta Giovanetti is funded by PON “Ricerca e Innovazione” 2014–2020 and by the CRP—ICGEB RESEARCH GRANT 2020 Project CRP/BRA20-03, contract CRP/20/03, Oswaldo Cruz Foundation. Stefano Pascarella is in part supported by the Sapienza grant n. RP12117A7670A1E8. The authors are grateful to the family of Prof. Giuseppe De Feo for the support. We also would like to thank all the authors who have kindly deposited and shared genomes on GSAID.

## Data Availability Statement

Genomes analysed in the present study were taken from GSAID database and are available at https://gisaid.org/.

## Author Contributions

*Conceptualization:* Fabio Scarpa, Daria Sanna, Marco Casu, Pier Luigi Fiori, Massimo Ciccozzi. *Data Analyses:* Fabio Scarpa, Stefano Pascarella. *Writing-original draft preparation:* Fabio Scarpa, Daria Sanna, Marco Casu. *Writing-review and editing:* Fabio Scarpa, Daria Sanna, Ilenia Azzena, Marco Casu, Piero Cossu, Pier Luigi Fiori, Domenico Benvenuto, Elena Imperia, Marta Giovanetti, Giancarlo Ceccarelli, Roberto Cauda, Antonio Cassone, Stefano Pascarella, Massimo Ciccozzi. *Resources:* Fabio Scarpa, Daria Sanna, Marco Casu.

